# Predicting the fundamental thermal niche of crop pests and diseases in a changing world: a case study on citrus greening

**DOI:** 10.1101/465898

**Authors:** Rachel A. Taylor, Sadie J. Ryan, Catherine A. Lippi, David G. Hall, Hossein A. Narouei-Khandan, Jason R. Rohr, Leah R. Johnson

## Abstract

1. Predicting where crop pests and diseases can occur, both now and in the future under different climate change scenarios, is a major challenge for crop management. One solution is to estimate the fundamental thermal niche of the pest/disease to indicate where establishment is possible. Here we develop methods for estimating and displaying the fundamental thermal niche of pests and pathogens and apply these methods to Huanglongbing (HLB), a vector-borne disease that is currently threatening the citrus industry worldwide.
2. We derive a suitability metric based on a mathematical model of HLB transmission between tree hosts and its vector *Diaphorina citri*, and incorporate the effect of temperature on vector traits using data from laboratory experiments performed at different temperatures. We validate the model using data on the historical range of HLB.
3. Our model predicts that transmission of HLB is possible between 16°C and 33°C with peak transmission at ~25°C. The greatest uncertainty in our suitability metric is associated with the mortality of the vectors at peak transmission, and fecundity at the edges of the thermal range, indicating that these parameters need further experimental work.
4. We produce global thermal niche maps by plotting how many months each location is suitable for establishment of the pest/disease. This analysis reveals that the highest suitability for HLB occurs near the equator in large citrus-producing regions, such as Brazil and South-East Asia. Within the northern hemisphere, the Iberian peninsula and California are HLB suitable for up to 7 months of the year and are free of HLB currently.
5. *Policy implications*. The thermal niche map indicates the places at greatest risk of HLB establishment should the disease enter these regions. This indicates where surveillance should be focused to prevent establishment. Our mechanistic method can be used to predict new areas for HLB transmission under different climate change scenarios and is easily adapted to other vector-borne diseases.

## 1 Introduction

The quality and quantity of yields for many crop systems can be significantly reduced by pests and disease. For example, the wheat aphid reduces yields of grain crops (Merrill *et al*., 2009), Pierce’s disease vectored by the glassy-winged sharpshooter diminishes grape yields (Galvez *et al*., 2010), codling moth damages apple orchards (Rafoss & Sæthre, 2003), and Huanglongbing vectored by the Asian citrus psyllid decimates citrus crops (da Graça *et al*., 2016). The pests and/or vectors of disease are often small arthropods that are sensitive to environmental conditions, including temperature and humidity (Mordecai *et al*., 2013; Tsai *et al*., 2002). Therefore, predicting when and where these pests or disease vectors will occur, and hence potential loss of crops, is often highly dependent on environmental conditions, and thus a changing climate. Some crops are already experiencing reduced yields associated with climate change (Challinor *et al*., 2014), an undesired outcome that could be exacerbated if it coincides with increases in pest populations or disease transmission (Cammell & Knight, 1992). However, predicting realistic impacts of climate change on living systems, such as pests and disease vectors, is a major challenge in ecology (Rohr *et al*., 2011).

With the pressing need to understand the effects of climate change on food production, we provide a method to estimate and display the fundamental thermal niche of a crop pest or disease. The fundamental thermal niche is the set of temperatures under which populations of a species would be expected to persist, all else being equal (Angilletta & Sears, 2011). It can be used to predict where pests or disease can currently establish outside of their existing range, as well as predict future locations of pest and disease outbreaks associated with a changing climate. This, in turn, can facilitate targeting risk-based surveillance and prophylactic interventions. Consequently, our method for estimating the fundamental thermal niche of a crop pest or disease should be an important tool for current and future agricultural planning.

In this study, we borrow approaches established for human vector-borne diseases (Mordecai *et al*., 2017, 2013) to simultaneously estimate the fundamental thermal niche of a crop pathogen and its insect vector. More specifically, we first develop a mechanistic model of disease transmission where the parameters of the model are fitted to data from temperature-dependent laboratory studies. A metric derived from this model is then used to indicate how suitable locations are based on average monthly temperatures. Finally, we validate the model by assessing how well it corresponds with observational data on known disease occurrence (Mordecai *et al*., 2013). Additionally, we use Bayesian inference to incorporate uncertainty arising from the use of multiple data sources and uncertainty in temperature dependence of each of the vector’s life history traits to estimate the overall uncertainty in the suitability metric (Johnson *et al*., 2015). Our method allows for the inclusion of the intrinsic reasons behind why an organism is found where it is and the potential interplay when different traits of organisms respond disparately to changes in extrinsic factors (Angilletta & Sears, 2011).

To develop our methods, we leverage data on the effect of temperature on bacteria of citrus that cause the disease Huanglongbing (HLB, or citrus greening) and their primary vector, the Asian Citrus Psyllid (*Diaphorina citri* Kuwayama; ACP). HLB is a devastating disease of citrus trees that has spread globally from its origin in Asia (Bové, 2006). It affects the quality and quantity of citrus fruit on a tree, for all citrus species, leading to misshapen fruit, bitter taste, and fruit dropping early (Bové, 2006). The symptoms can be difficult to detect, and may take months to appear on a tree, but include chlorosis of leaves with eventual dieback and death of the tree (Gottwald, 2010; Lee *et al*., 2015). HLB is caused by three bacteria: *Candidatus* Liberibacter asiaticus (CLas), *Candidatus* Liberibacter africanus (CLaf), and *Candidatus* Liberibacter americanus (CLam) (Bové, 2006). The predominant bacterium, and the focus in this study, is CLas, which occurs in all HLB-infected areas (Gottwald, 2010) apart from South Africa (where CLaf is present). CLam occurs primarily in South America alongside CLas, and is responsible for only a minority of cases there (Gottwald, 2010). Transmission of both the CLas and CLam bacteria occurs due to feeding of the ACP (Grafton-Cardwell *et al*., 2013; Hall *et al*., 2013). ACP and HLB have spread throughout the world mostly via worldwide trade (Byrne *et al*., 2018), and now exist in nearly all citrus producing regions (Hall *et al*., 2013). The cost of this disease to the citrus industry is huge, and interventions to prevent its spread and reduce the deleterious effects of the disease are, for the most part, ineffective (Hodges & Spreen, 2012). Here, we map the suitability metric of HLB around the world to provide a planning tool for citrus growers, whilst the method itself is applicable to all crop diseases spread by vectors or to crop pests.

## 2 Methods

### 2.1 Formulation of *S*(*T*)

To determine the thermal niche of citrus greening, we characterize the possibility of an introduction of citrus greening persisting at a local level, for different locations around the world. We use an estimate of the basic reproductive number *R*_0_. When *R*_0_ > 1, the disease is likely to spread and lead to an epidemic, whereas if *R*_0_ *<* 1, the disease will die out. A previously developed model for the dynamics of citrus greening between trees and psyllids within a single grove is used to determine *R*_0_ (Taylor *et al*., 2016). In this mechanistic model, trees and psyllids are split into different compartments based on their disease status. Trees are Susceptible, Asymptomatic (infected, but no symptoms) or Infected (with symptoms). Psyllids are Susceptible, Exposed or Infected (Fig 1). Death of trees and roguing (removing trees from a grove due to high levels of infection) are included, as well as replacement of all removed trees with susceptible trees. For full details of this model, see Supplementary Info S1. The basic reproductive number *R*_0_ is then calculated as (Taylor *et al*., 2016):

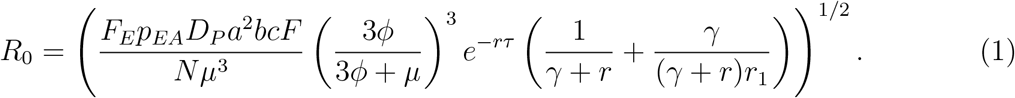

**Figure 1:**
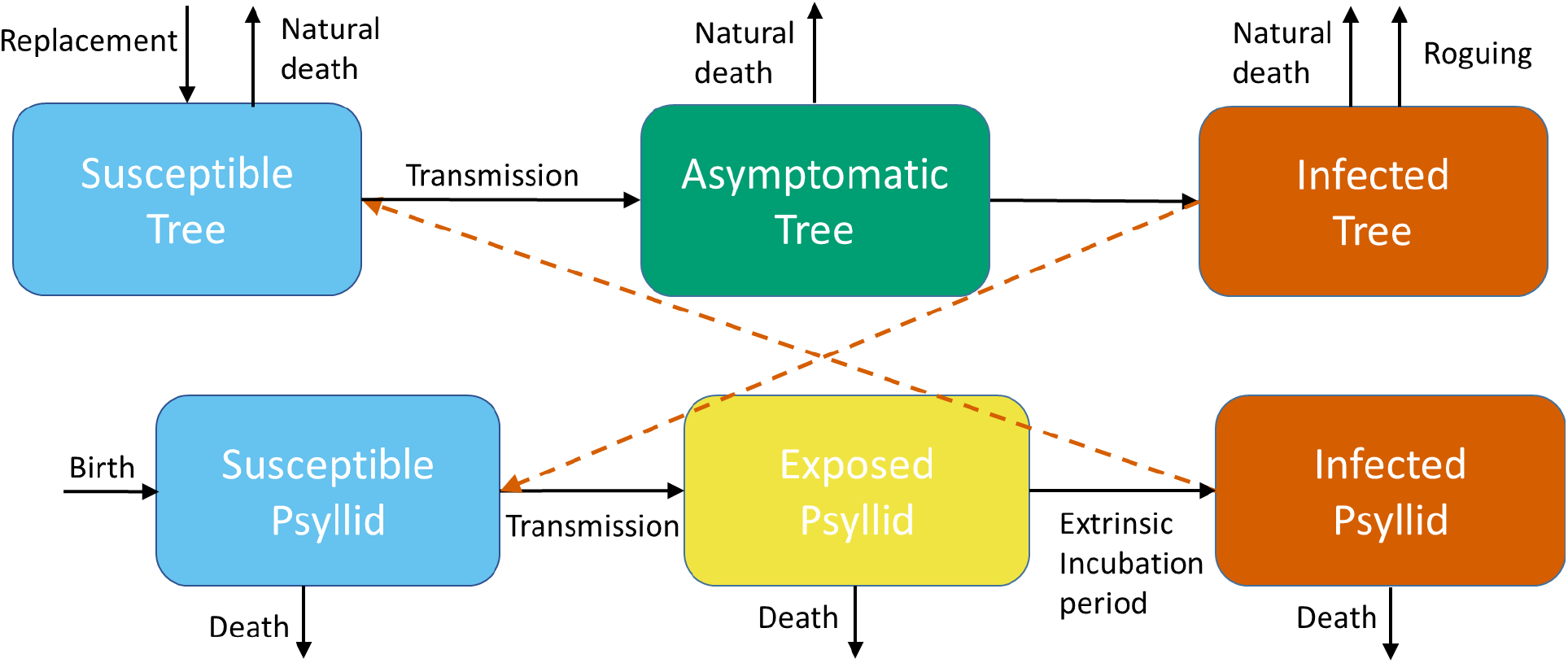
A schematic of the model for HLB transmitted between trees and psyllids. Trees are Susceptible, Asymptomatic, or Infected. Psyllids are Susceptible, Exposed or Infected. Dead and rogued trees are replaced by susceptible trees. Black arrows show the transitions between the compartments. Orange dashed arrows show the necessary interactions between trees and psyllids to obtain transmission.

This equation for *R*_0_ can be understood by considering how disease propagates through the citrus system. The number of psyllids in the population is determined by:

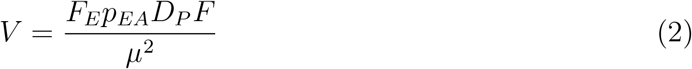

which includes the fecundity of adult psyllids (*F_E_*), the probability of egg to adult survival (*p_EA_*), the development rate from egg to adult (*D_P_*), the mortality rate of adult psyllids (*µ*) and the amount of trees flushing (*F*) within the grove which varies throughout the year. These adult psyllids are in contact with the single infected tree based on the bite rate (*a*) with a probability of transmission from tree to psyllid (*c*). The psyllids undergo an extrinsic incubation period before becoming infectious given by rate (*ϕ*). But they can die during this time which leads to the term 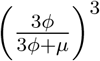for the number of psyllids that survive the extrinsic incubation period. These infectious psyllids are in contact with susceptible trees (total *N*), once again with bite rate *a* and a probability of transmission from psyllid to tree (*b*). The term *e^−rτ^* represents the proportion of trees that survive the incubation period (*τ*) to become infectious. The final combined term in the equation determines how long a tree is infectious, during both the asymptomatic stage and the infected stage, and includes the rate at which trees develop symptoms (γ), the death rate of asymptomatic trees (*r*) and the death rate of infected trees (*r*_1_).

We define our measure of thermal suitability for HLB as the vector/infection components of *R*_0_ that depend on temperature only. That is, the suitability, *S*(*T*), is given by:

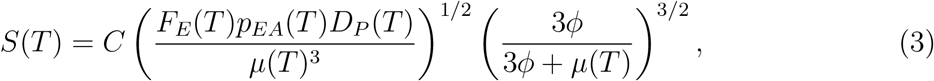

where *C* is a constant that scales the mean suitability to lie between 0 and 1. Thus, the suitability is zero when temperature is predicted to fully exclude transmission and 1 at maximal transmission. In the results, we will primarily focus on predictions based on two suitability regimes: a “permissive” thermal niche corresponding to temperatures where *S*(*T*) > 0; and a “highly suitable” thermal niche corresponding to temperatures such that *S*(*T*) > 0.75 (i.e., the highest quartile of suitability).

### 2.2 Bayesian fitting of thermal traits in *S*(*T*)

It has been widely recognized that performance traits of ectotherms, such as survival, reproduction, and movement, exhibit unimodal responses to temperature (Amarasekare & Savage, 2012; Dell *et al*., 2011). Following the approach introduced in Mordecai *et al*. (2013), we fit unimodal temperature responses to laboratory data for traits of the vector that appear in the equation for *S*(*T*). These curves can then be inserted into the equation for *S*(*T*) to determine how transmission depends upon temperature. As in Johnson *et al*. (2015), we take a Bayesian approach to fitting. Thus, we can quantify the uncertainty in the temperature relationship of individual components and explore the emergent uncertainty in *S*(*T*) overall that is due to the uncertainty in the components. Complete details of the approach are presented in Supplementary Info S2.

Here we focus on four psyllid traits for which there exists data across sufficient temperatures to quantify the responses: fecundity (*F_E_*); probability of egg to adult survival (*p_EA_*); average longevity (the inverse of the mortality rate, 1*/µ*); and the development rate of psyllids from eggs into adults (*D_P_*). The majority of the data comes from Liu & Tsai (2000), which has a range of 15°C up to 30°C for all four parameters, with additional data at 10°C and 33°C for some of the parameters. Hall *et al*. (2011) and Hall & Hentz (2014) provide data on high and low extreme temperatures that prevent development of the psyllids and/or lead to mortality. However, Hall *et al*. (2011) also provide information on fecundity of female psyllids across a temperature range of 11°C to 41°C. We include both data sets for fecundity, and consider whether they generate different predictions. Hereafter, Liu & Tsai (2000) and Hall *et al*. (2011) will be referred to as LT00 and H11 respectively and, when referring to the *S*(*T*) output created by either data set, LT00 *S*(*T*) and H11 *S*(*T*) will be used.

For all sets of data, we fit two functions to describe the mean relationship between the trait and temperature: quadratic, giving a symmetric relationship (*f*(*T*) = *qn*(*T* − *T*_0_)(*T* − *T_M_*)); Brière, giving an asymmetric relationship (*f*(*T*) = *cT* (*T* − *T*_0_)(*T* − *T_M_*)^1/2^, Brière *et al*. (1999)). All responses were fitted using a Bayesian approach. After specifying the mean relationship, probability distributions appropriate to describe the variability around this mean were chosen, and priors specified (Supplementary Info S2). We chose priors to limit parameters for the minimum and maximum temperature thresholds to approximate known limits to psyllid survival. For instance, the prior on the thermal limits was set uniformly over the interval from 30°C-50°C to acknowledge that exposing psyllids to the very high temperatures kills them almost immediately, while being wide enough to allow the laboratory data primacy in the analysis. Priors for other parameters of the mean were chosen to be relatively uninformative but scaled appropriately for the response. Priors on variance parameters were typically also chosen to be relatively uninformative, although the precise specification varied due to differences in the scale of the responses and to improve mixing and convergence of the Markov Chain Monte Carlo sampling scheme. All responses were fitted in R (R Development Core Team, 2008) using the `JAGS/rjags` packages (Plummer, 2003, 2013). After fitting both quadratic and Briére responses to each data set, the preferred response was chosen via Deviance Information Criterion (DIC) as implemented in `rjags`.

Once samples of the posterior distributions of parameters for the preferred model were obtained, these were used to calculate the posterior samples of the response across temperature. Then, at each temperature the mean/median of the response and the 95% highest posterior density (HPD) interval were calculated. The posterior samples of each trait response were then combined to create samples from the overall response of *S*(*T*) to temperature. As with the individual thermal responses, these posterior samples of *S*(*T*) across temperature were used to calculate the mean and 95% HPD interval around the mean of *S*(*T*).

### 2.3 Uncertainty in response of *S*(*T*) to temperature

The uncertainty for each parameter in *S*(*T*) is calculated using the variation in *S*(*T*) at each temperature when all parameters, apart from the one of interest, are held constant at their mean posterior values for that temperature. We calculate the 2.5 and 97.5 quantiles of the *S*(*T_M_*) posterior distribution and plot the difference against *T_M_*. We do this for values of *T_M_* in our temperature range, and then for all parameters in *S*(*T*). This method indicates which parameters have the greatest variation at each temperature and hence which parameters have the greatest uncertainty at that temperature. This allows us to know when our estimate of *S*(*T*) is uncertain and which parameter is causing this.

### 2.4 Validation of suitability measure

Narouei-Khandan *et al*. (2016) present spatially explicit data on locations with confirmed presence of the Asian citrus psyllid (ACP) or HLB (CLas form specifically). These are presence-only data, so we cannot examine how well our model partitions predictions between suitable and unsuitable areas. Instead, we focus on a more qualitative assessment of model adequacy. For each location in the dataset, we calculate the number of months that the average temperature falls within the bounds of the permissive or the highly suitable thermal ranges of ACP from our model. If our model can adequately capture temperature conditions related to transmission and vector establishment, then most locations of HLB presence will have many months in the suitable range.

### 2.5 Mapping suitability across space

To communicate the potential suitability of the world’s land surface for transmission of HLB, we mapped the months of suitability as a function of mean monthly temperatures. We demonstrate two subsets of suitability, the most optimal (“highly suitable”) and the overall suitability (“permissive”), namely the top 25th percentile and 100th percentile of the *S*(*T*) curve (Ryan *et al*., 2015). To derive these percentiles, we took the posterior mean *S*(*T*) curves for the H11 *S*(*T*) and LT00 *S*(*T*) models across temperatures and extracted the temperatures corresponding to the top 25th percentile of the *S*(*T*) curve (i.e., *S*(*T*) > 0.75, Supplementary Info S5), the highly suitable thermal niche, as well as the temperatures corresponding to the transmission limits (*S*(*T*) > 0), corresponding to the permissive thermal niche. These values were used with the climate models (§2.5.1) to determine, on average, the number of months each year that each pixel is either permissive or highly suitable.

### 2.5.1 Climate models

We mapped the suitability measures onto rasters of current mean monthly temperature data at 0.1°C intervals. Data were derived from WorldClim Version 1.4 dataset (www.worldclim.org) (Hijmans *et al*., 2005) at 5 minute resolution (roughly 10km^2^ at the equator). The scaled suitability model was projected onto the climate data using the ‘raster’ package (Hijmans, 2016) in R (R Development Core Team, 2008). R code for this step is supplied in the Supplementary Material. Visualizations were generated in ArcMap. For each of the scenarios, we created global maps and insets for areas of citrus growing concern in the Northern Hemisphere: California, Florida, and the Iberian peninsula.

## 3 Results

### 3.1 Posterior distributions of thermal traits

The probability of egg to adult survival (*p_EA_*) and longevity (1*/µ*) are both fitted best by quadratic curves, while development rate from egg to adult (*D_P_*) is best fitted by a Briére curve (Fig 2). However, fecundity (*F_E_*) switches from a Briére to a quadratic depending on whether we use the data from Liu & Tsai (2000) or Hall *et al*. (2011) respectively (Fig 2E,F). The data sources also predict different upper thermal limits for fecundity: LT00 predicts no fecundity above 31°C, whereas fecundity is possible up to 41°C according to H11. Full posterior plots of the parameters are in Supplementary Info S3.

**Figure 2:**
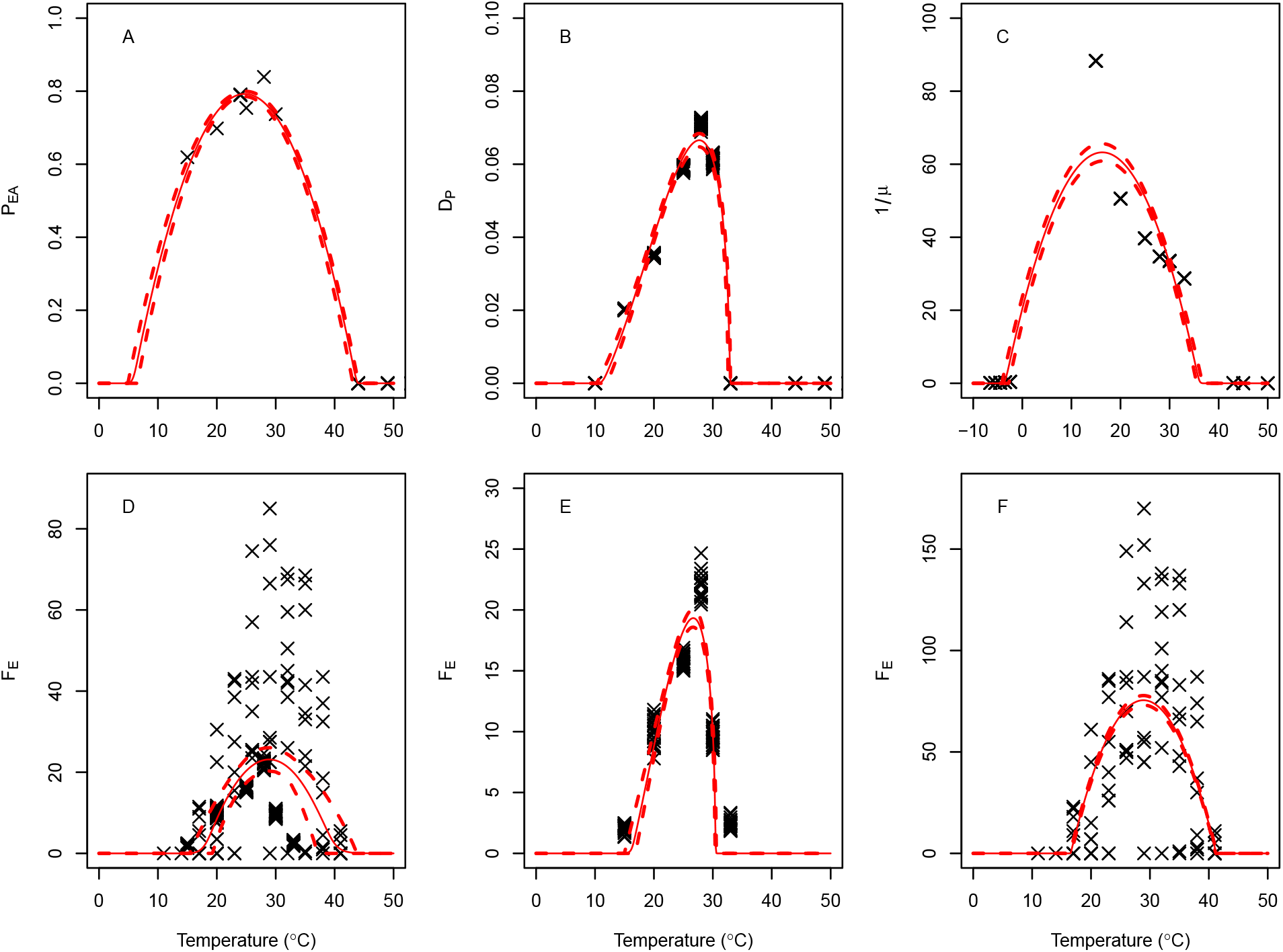
Psyllid trait data against temperature (°C) with the best fit plotted as a solid line and 95% quantiles as dashed lines. In *A*, the probability of egg to adult survival (*p_EA_*); in *B*, the development rate from egg to adult psyllid (*D_P_*); in *C*, the longevity of adult psyllids (1*/µ*); in *D − F*, the fecundity of adult psyllids (*F_E_*). In *D*, both data sets are plotted together, whereas in *E*, only the Liu & Tsai (2000) data is used and in *F*, only the Hall *et al*. (2011) data is used.

### 3.2 Posterior distribution of *S*(*T*)

The lower thermal bound of the posterior distributions of LT00 and H11 *S*(*T*) are in agreement, predicted using the two different data sets for fecundity, although LT00 *S*(*T*) has more uncertainty (Fig 3, Supplementary Info S4). The temperature at which the peak of *S*(*T*) occurs is also very closely aligned. However, the value of *S*(*T*) at that peak temperature and the upper thermal limit are very different. When scaled so that the LT00 *S*(*T*) has a maximum of 1, the peak of H11 *S*(*T*) is 1.35 times larger. LT00 *S*(*T*) is driven to 0 at approximately 31°C because fecundity is not possible for higher temperatures. However, H11 *S*(*T*) still has transmission predicted up to 33°C.

**Figure 3:**
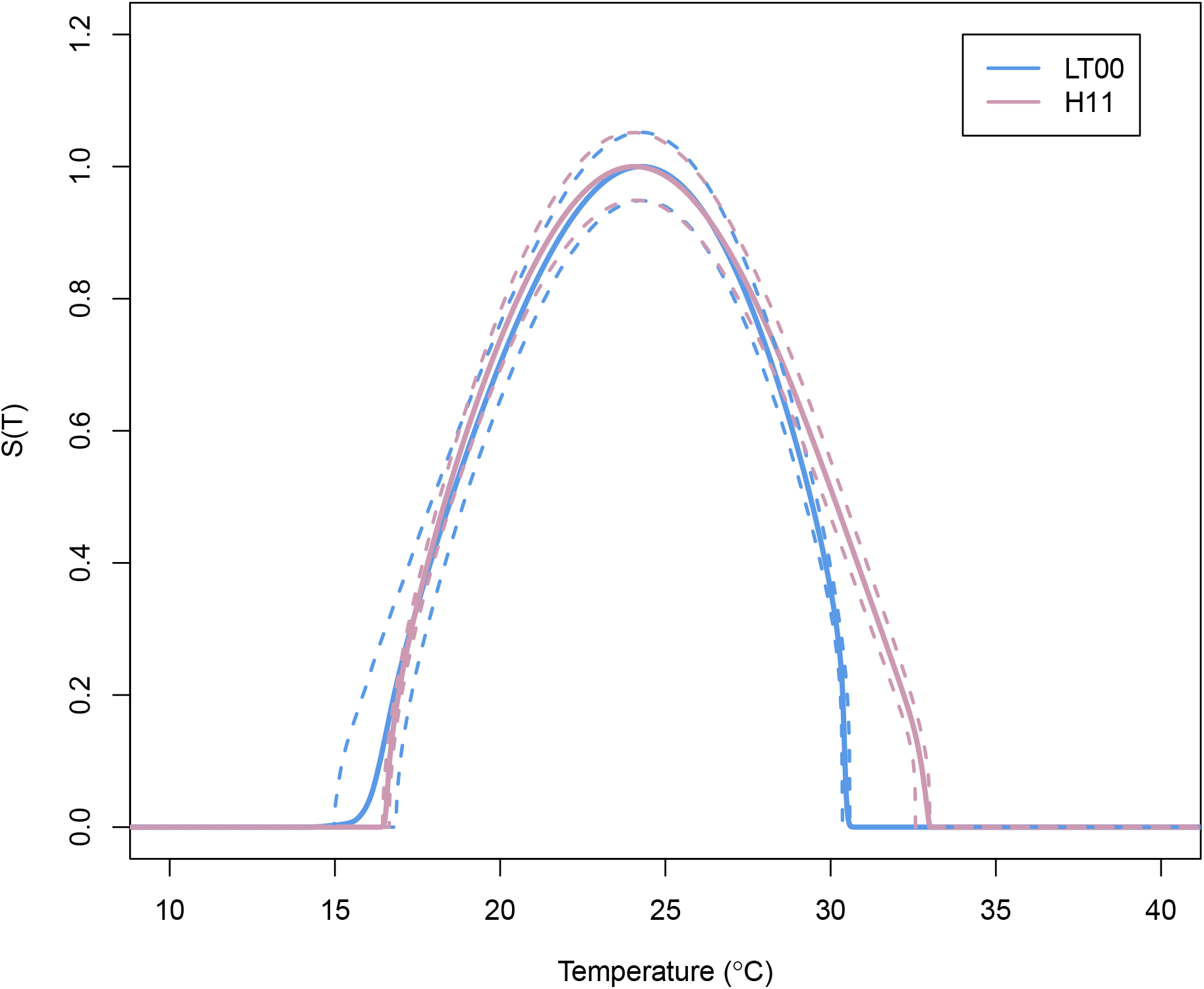
Posterior distribution of *S*(*T*) against temperature (°C) using data from Liu & Tsai (2000) (LT00, in blue) and Hall *et al*. (2011) (H11, in pink). Mean *S*(*T*) for both models is plotted using solid lines, 95% credible intervals are plotted with dashed lines.

### 3.3 Sources of uncertainty in *S*(*T*)

The uncertainty of each parameter on *S*(*T*) is plotted against temperature as all other parameters are held constant at their means (Fig 4). We can use Fig 4 to understand what drives *S*(*T*) at different temperatures, and therefore it indicates how best to affect *S*(*T*) at those temperatures, if the aim is intervention. In Fig 4*A*, fecundity (*F_E_*) is the main parameter driving variability in *S*(*T*) during the range 15-20°C as well as when *S*(*T*) is decreasing to 0 at 31°C. However, adult mortality (*µ*) is the main proponent of uncertainty during the mid to high temperatures of 20-30°C. In comparison, in Fig 4, adult mortality (*µ*) leads to the most uncertainty over the whole temperature range. While adult fecundity is once again important at low temperatures, it is the development rate (*D_P_*) that emerges as producing the most variability in *S*(*T*) at high temperatures.

**Figure 4:**
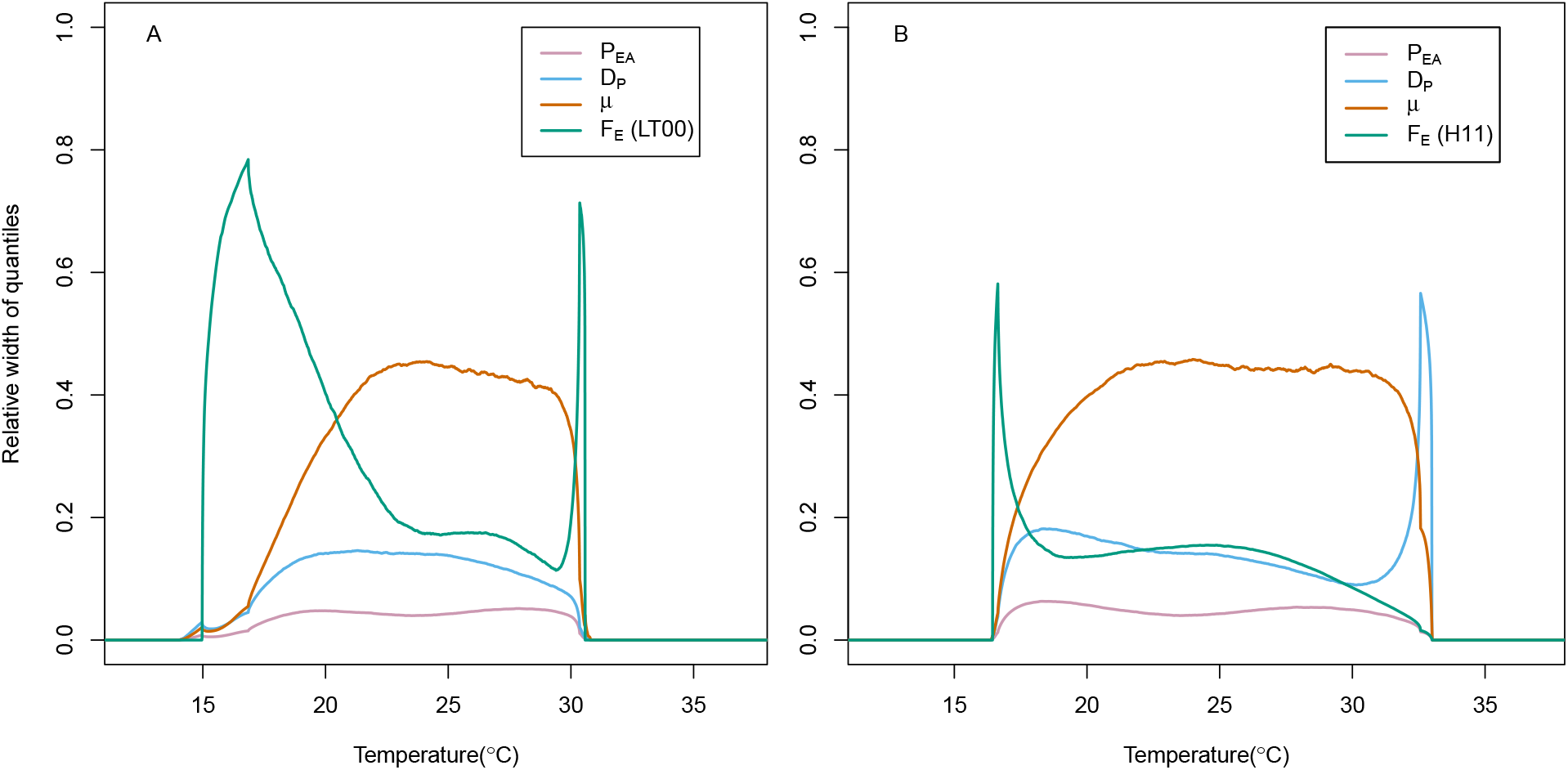
The uncertainty of *S*(*T*) propagated through each parameters’ relation to temperature. In (*A*) uncertainty in LT00 *S*(*T*) and in (*B*) uncertainty in H11 *S*(*T*). This is produced based upon the posterior of *S*(*T*), holding all parameters constant at their predictive mean apart from the parameter of interest. The 2.5% and 97.5% quantiles of the resultant estimation of *S*(*T*) are then calculated at each temperature and the difference between these quantiles is plotted.

### 3.4 Validation

We present histograms of the number of months that the average temperature falls within the bounds of both the permissive or the highly suitable thermal ranges (Fig 5, based on the H11 model) at each location in the data set from Narouei-Khandan *et al*. (2016). We restrict ourselves to locations from that dataset that are not from mountainous regions (i.e. exclusion of ACP(n=26) and HLB(n=12) points recorded at relatively high elevations, coinciding with named mountain ranges). This is because, at the spatial scale we use, these areas tend to have much more variation in temperature resulting in higher uncertainty in the temperature predictor than in other locations.

**Figure 5:**
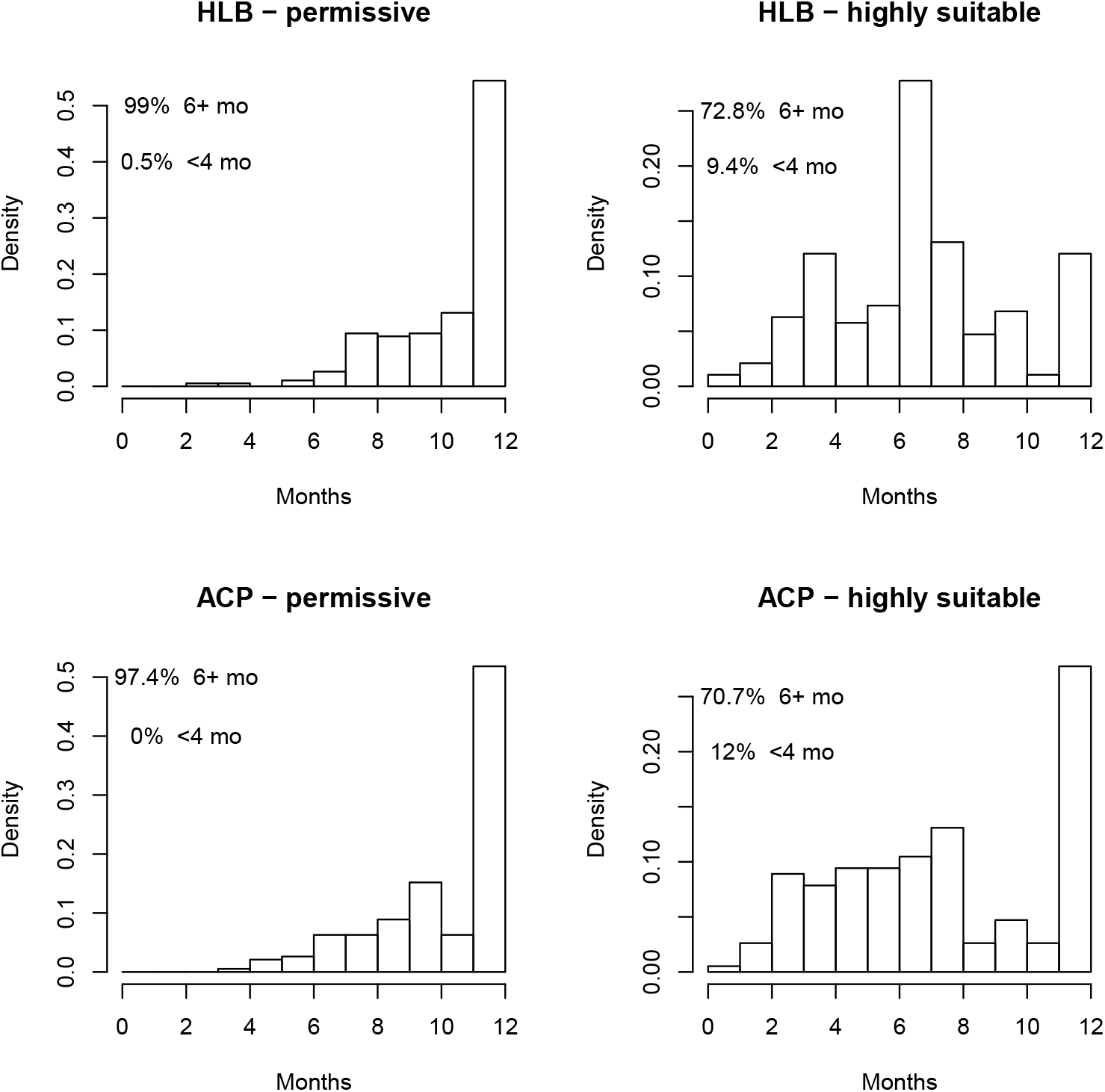
The number of months that every location with current HLB or ACP presence is either permissive or highly suitable according to our H11 *S*(*T*) model. Top row: locations in the validation dataset where HLB is present. Bottom row: locations in the validation dataset where ACP is present. We define permissive suitability as *S*(*T*) > 0 and high suitability as *S*(*T*) > 0.75.

Most locations with HLB or ACP have permissive temperature ranges for at least six months of the year. Over 82% of locations are in the permissive range for nine or more months of the year, and more than 50% have permissive temperatures year round. Locations with year round highly suitable conditions account for 12% and 28% of records for HLB and ACP, respectively, and over 70% of locations have highly suitable conditions for 6 months or more. There are almost no locations with HLB or ACP present that the conditions are not permissive for 3 months or more. Results based on the data by Liu & Tsai (2000) are similar, as are the results that include mountainous areas (see Supplementary Info S6)

### 3.5 Thermal niche of HLB

In Fig 6 and Fig 7, we present the mapped outputs of the permissive and highly suitable regions, respectively, for H11 *S*(*T*). Similar maps for the LT00 *S*(*T*) model are presented in Supplementary Info S7. Many locations in the Southern hemisphere are permissive for HLB all months of the year (Fig 6), including in South America and Southwest Asia where the disease is currently present. Australia and many countries in Africa, which are large citrus producing regions, are permissive for HLB all or many months of the year, but CLas HLB is not currently present. The insets highlight that southern Florida is permissive for HLB all months of the year, and for at least 7 months in the north. This is confirmed on the ground as HLB is present throughout the whole state of Florida. California and the Iberian peninsula have similar suitability profiles to each other, with up to 7 months permissive in the south of the Iberian peninsula, and up to 8 months at the very south of California. As expected, more northerly regions of the world, which are not suitable for growing citrus, are also not able to maintain ACP populations.

**Figure 6:**
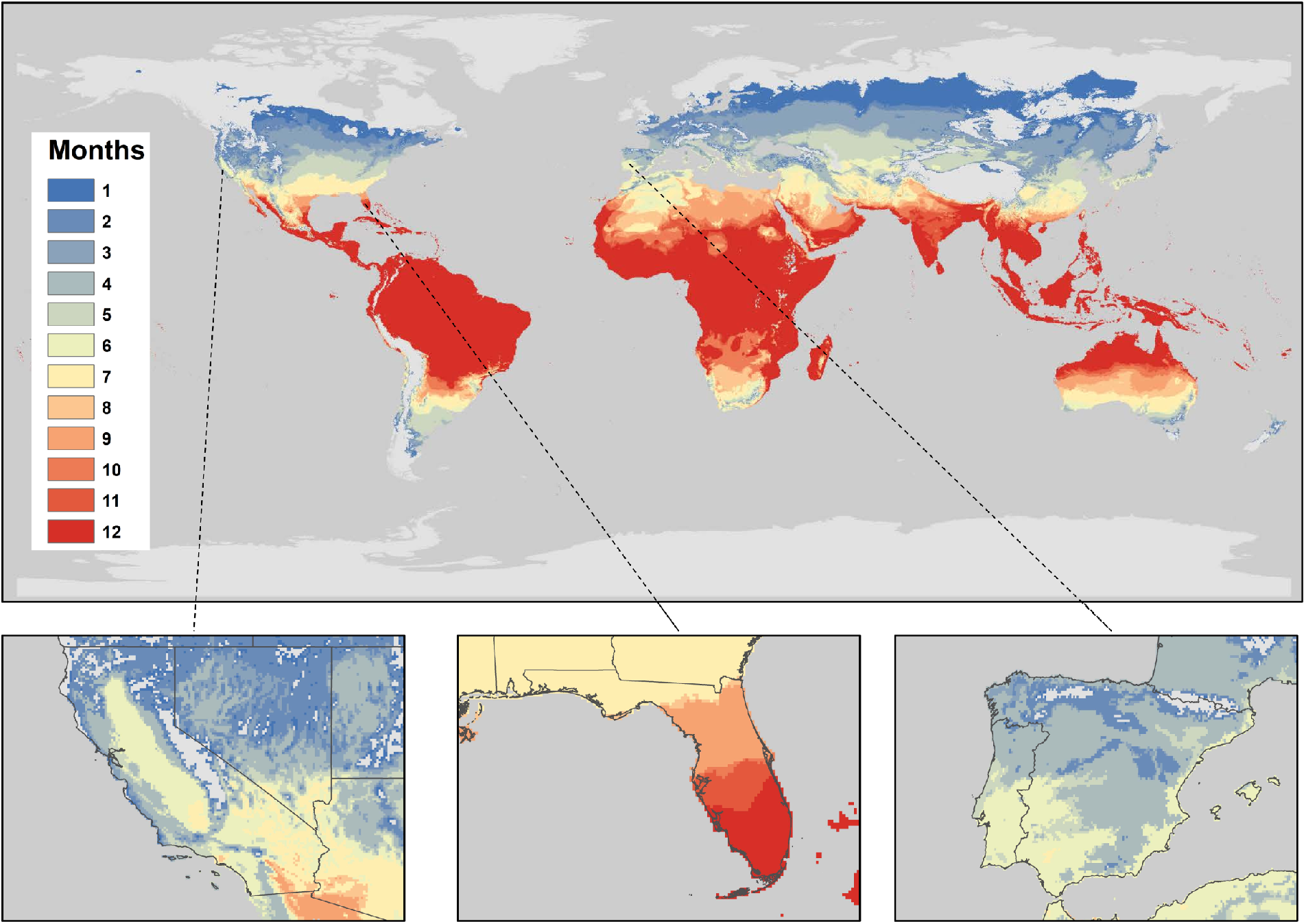
The number of months a year that locations have permissive temperatures according to our H11 *S*(*T*) model. Inset plots of California, Florida and the Iberian peninsula, respectively, are included. We define permissive temperatures for suitability as *S*(*T*) > 0. Locations in grey have zero months suitable for HLB transmission.

**Figure 7:**
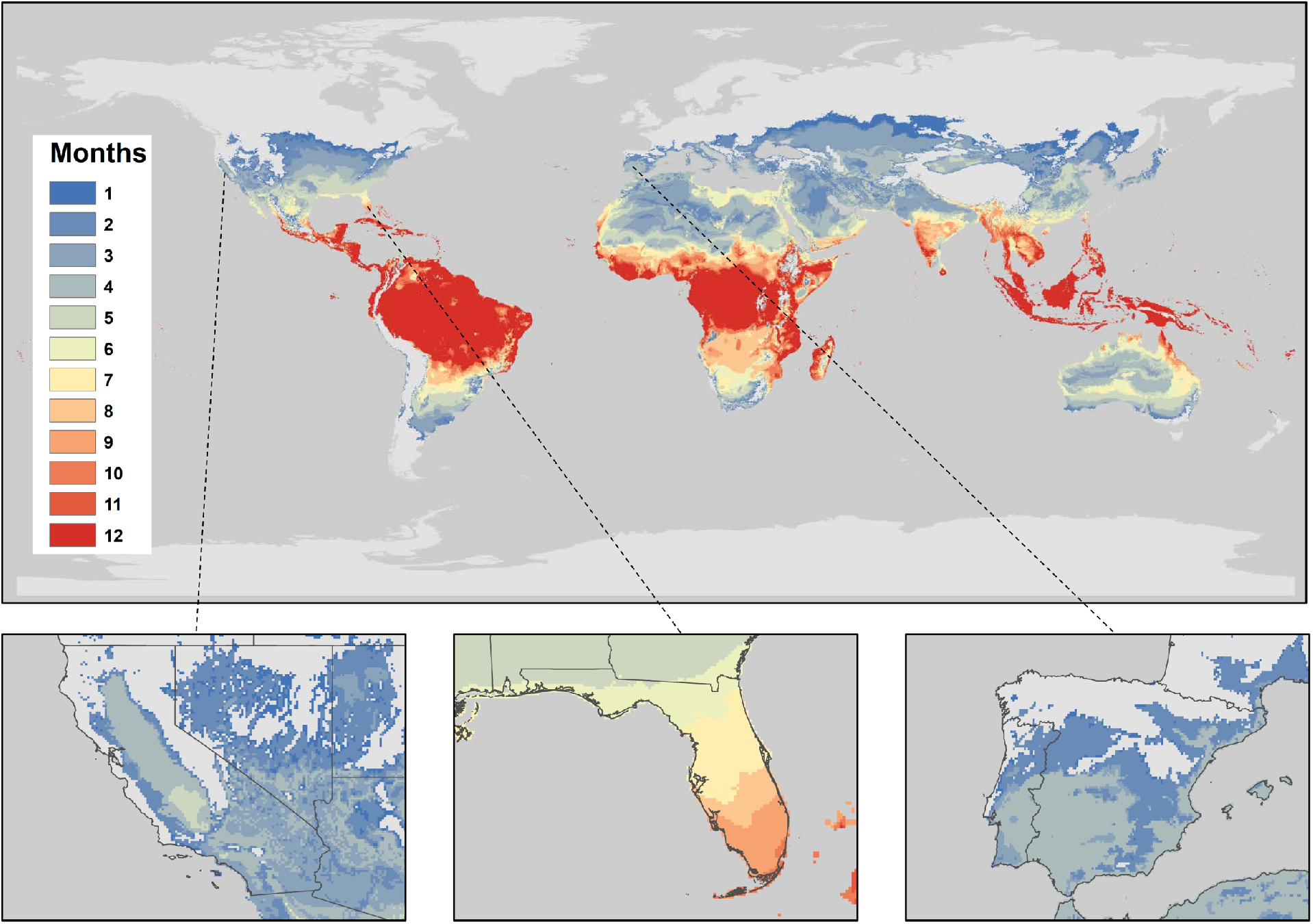
The number of months a year that locations have highly suitable temperatures according to our H11 *S*(*T*) model. Inset plots of California, Florida and the Iberian peninsula, respectively, are included. We define highly suitable temperatures as *S*(*T*) > 0.75. Locations in grey have zero months suitable for HLB transmission.

The pattern for suitability is similar for the highly suitable map (Fig 7) but with lower numbers of months satisfying this stricter criteria. South America maintains year round high suitability for the disease across much of the continent. Similarly, Southwest Asia maintains suitability year-round, whereas Australia is reduced to 7 months or less across the country. California and the Iberian peninsula are highly suitable for 4-5 months of the year, whereas Florida is still highly suitable for up to 9 months each year.

## 4 Discussion

In this paper we presented a model of suitability for a specific crop disease, HLB. This method of demonstrating duration of risk of potential disease emergence and transmission, as a function of thermal suitability, has been successfully used for human vector-borne diseases such as dengue, Zika, and malaria (Mordecai *et al*., 2017, 2013). This climate-predictive mapping framework provides a tool for planning and intervention, and is adaptable to multiple systems, including vector-borne diseases of crops and thermally sensitive crop pests. While we have demonstrated this approach for a coupled vector-pathogen system, and thus used *R*_0_ as a starting point to create a transmission-based suitability metric, the method to create a fundamental thermal niche for invasive crop pests is readily possible using population models (in which *R*_0_ represents the likelihood of population persistence rather than disease persistence, or for instance using *r* the intrinsic rate of population increase as an alternative measure). Given the availability of rigorous laboratory experiments on the thermal responses of other crop pest or disease systems, this approach is broadly applicable to many systems.

Our model outputs a suitability metric *S*(*T*) for transmission of HLB dependent on temperature. Conditional on the data we use to parameterize *S*(*T*), we predict that transmission can occur between 16°C and 30-33°C with peak transmission at approximately 25°C. While the lower bound and peak temperature predictions are similar regardless of which of the two fecundity data sets we use for parameterization, i.e., those from Hall *et al*. (2011) (H11) or Liu & Tsai (2000) (LT00), the predicted upper limit of suitability of the peaks differ depending on the data used (Fig 3). More specifically, the range is wider for the H11 data. Thus, based on the suitability index, there might be more areas strictly suitable for transmission if we assume the H11 data are more representative of psyllid populations in the field. Furthermore, when we consider our uncertainty analysis, there are additional differences between the two data sources (Fig 4). The main uncertainty in the LT00 model arises from fecundity at lower temperatures and mortality for higher temperatures, whereas for the H11 model, mortality is the main driver of uncertainty overall. In both cases, mortality of ACP is the most important parameter when *S*(*T*) is near its peak at 25°C. Together, these indicate that focus should be on further experimental understanding of mortality rates and fecundity near the edges of the thermal tolerances of psyllids to refine our estimates of the thermal niche.

We use our suitability metric to predict regions which are permissive or highly suitable for HLB around the world. Our maps indicate that regions close to the equator have the greatest number of months permissive for HLB transmission, especially in South America, Africa and South East Asia. South America and South East Asia are, in particular, large citrus producing regions, and HLB is already present in both (Coletta-Filho *et al*., 2004; Garnier & Bové, 1996, 2000; Torres-Pacheco *et al*., 2013). The fact that HLB is not only permissive all 12 months of the year, but is actually highly suitable year-round, indicates the scale of the potential problem in those areas. However, the HLB epidemic in São Paulo State in Brazil, the major citrus producing region in the country, is successfully managed in those areas that have adhered to strict recommendations for control (Belasque *et al*., 2010). The disease entered the state in 2004 (Coletta-Filho *et al*., 2004). By 2012, incidence of disease was estimated as low as 1% for symptomatic trees across a third of the citrus acreage in the state (Bové, 2012). In comparison, in Florida, where the disease was first discovered in 2005, growers in a survey in 2015 were asked to estimate both the percentage of their citrus acres with at least one tree infected and the percentage of all their citrus trees infected, with results indicating 90% and 80%, respectively (Singerman & Useche, 2016). This is despite the fact that our suitability metric estimates the northern regions of Florida to be highly suitable for HLB only 6 months of the year. While Brazil might have managed to control HLB successfully in some regions, the disease is still spreading throughout the country in those regions which have not been as proactive in their control (Belasque *et al*., 2010); a reminder that, without strict control measures, the disease can spread quickly and devastatingly, with 100% HLB incidence possible in infected groves (Bové, 2012).

Within the northern hemisphere, we highlight the suitability of California and the Iberian peninsula for HLB transmission. Although California has had incursions of the disease, with the first occurring in 2012 (Kumagai *et al*., 2013), all have been in trees at residential properties and hence the citrus industry in California is currently free from disease (Byrne *et al*., 2018). Similarly, the Iberian peninsula has had no cases of HLB (Cocuzza *et al*., 2017). However, both regions have high citrus production and thus the potential consequences of incursion of HLB are high. They have similar suitability profiles with permissive suitability of HLB on average about 6 months of the year in both regions. While disease transmission might not be permissible all year round, trees can remain infected with HLB indefinitely unless they are rogued, thus allowing over-wintering of the disease during the seasons that ACP will not be active (Gottwald, 2010). Therefore, for these two regions to maintain HLB-free status, they need to deal with incursions promptly to ensure infected trees are removed. However, California has the added disadvantage that its neighboring country and many neighboring states have the disease or have ACP present (Torres-Pacheco *et al*., 2013). Indeed, the initial incursion of ACP into California is most likely to have occurred from Mexico (Bayles *et al*., 2017). This makes it harder to reduce the likelihood of HLB incursion as it is difficult to control the movement of vectors across borders. For the Iberian peninsula, it has been suggested that incursion of HLB is most likely through contaminated trade, such as infected plant materials (Cocuzza *et al*., 2017).

Our analysis has been performed for the transmission of the CLas form of HLB by the vector ACP and thus does not quantify the spread of CLaf HLB around the world from the African citrus psyllid (AfCP). Therefore, the suitability for HLB transmission might be underestimated in some areas of the world since AfCP has a different temperature profile than ACP, as it is more sensitive to heat (Schwarz & Green, 1972) and the two psyllids have not been found in the same locations (da Graça *et al*., 2016). In 2014, AfCP was first discovered in mainland Spain, alarming the citrus industry in Spain and Portugal (Cocuzza *et al*., 2017). Thus, it is possible that the Iberian peninsula is permissive for HLB for more months of the year than we have predicted if AfCP is the primary vector there.

We have validated this model using spatially explicit records of HLB and ACP presence. Most areas with confirmed HLB or ACP are in regions our model predicts as permissive or highly suitable for most of the year, indicating that our temperature-only model can capture an important component of the environment that constrains the spatial distribution of HLB. Narouei-Khandan *et al*. (2016) estimate from their HLB niche model that annual precipitation levels are the greatest predictor of HLB presence around the world. Unfortunately, there are not enough laboratory experiments assessing the effect of humidity on psyllid life history traits for us to include this in our model. Whilst our validation results indicate that we have successfully characterized a significant component of HLB transmission using temperature, our predictions could undoubtedly improve if we also included the effects of humidity. Our model produces similar results to the model of Narouei-Khandan *et al*. (2016), although it is difficult to make comparisons between number of permissive months (our model) versus the probability of occurrence (Narouei-Khandan *et al*. (2016) model). The Narouei-Khandan *et al*. (2016) model predicts greater areas in Australia suitable for HLB and ACP establishment than our model but only coastal areas of California are predicted to have a high probability of HLP and ACP occurrence with a low probability elsewhere in California. In contrast, our model predicts up to 7 months of permissive suitability across much of California.

Our method for creating the thermal niche is adaptable to other crop diseases and pests due to its strength of being built using mechanistic models. Spatially-explicit data of disease presence are typically used to build ecological niche models in other contexts. A mechanistic model with additional on-the-ground validation is likely to predict more robustly how temperature constrains transmission than a correlative approach based on presence-only data. It also enables clean projections for future climate scenarios, as it is not limited by unquantifiable changes in land cover.

Our use of two data sets for one parameter highlights how our understanding of a disease/pest population can change significantly depending on the data we use to parameterize our models. The best way to avoid this and reduce our uncertainty is to use multiple data sources combined, but this requires multiple experiments from different laboratories estimating the same parameter across a range of temperatures using the same empirical approaches. Often multiple experiments like this do not take place because of a perceived lack of novelty, but as we show here, they are potentially important to fully understand the impact of temperature on the persistence and establishment of vector-borne diseases and pest populations.

As our mapping of suitability is performed at the pixel level, approximately 10km^2^ at the equator, this allows us to predict suitability to a very fine scale. Therefore, we can use the map as a tool to determine surveillance and management strategies for HLB as well as using the method for other vector-borne crop diseases or pests. Suitability does not indicate where incursions of the disease are likely to occur, but it does highlight the regions where it is most likely to establish and therefore where it is most necessary to avoid incursion. Thus, surveillance should be higher in the more suitable regions. For example, for HLB in California, surveillance should be targeted mostly towards the southern part of the state, and for Australia, towards the northern portion of the country (Fig 6). Furthermore, for those countries with disease present already, the suitability maps can indicate which regions should have different management aims. In areas with very high suitability, a strict management policy might aim to keep incidence levels low. On the other hand, the aim might be complete eradication in regions with lower suitability. Or the management strategy might be implemented only during those months suitable for the disease vector/pest. A more focused surveillance and management strategy produced with suitability maps can save time, money and resources, which can be valuable considering the economic costs currently involved in managing crop diseases (Challinor *et al*., 2014). São Paulo State, Brazil, demonstrates that it is possible to keep incidence of HLB low, even in a region which is highly suitable for HLB all 12 months of the year. Vera-Villagrán *et al*. (2016) estimated the economic benefits of implementing a Brazilian strategy in Mexico and found that it was cost-effective, assuming all growers abide by the regulations. The costs of implementing such a strict control strategy may be prohibitive, but it gives hope for the industry that control is possible, especially if implemented as soon as HLB is discovered (Belasque *et al*., 2010). Similarly, for other vector-borne diseases of crops and crop pests, successful management and control are possible if implemented quickly and extensively after disease or pest emergence. Overall, our suitability maps provide an additional tool, alongside modeling of intervention strategies, cost-benefit analysis, experimental studies, development of disease-resistant trees and other inventions, in the fight against vector-borne crop diseases and pests.

## Author Contributions

RAT, SJR, JRR and LRJ conceived and designed the study. RAT created the model and performed the Bayesian analysis. CAL and LRJ performed the validation, SJR performed the spatial mapping. HAN and DGH provided data. RAT, SJR and LRJ wrote the manuscript. All authors reviewed the manuscript and gave approval for publication.

## Acknowledgements

LRJ and JRR were supported by the National Science Foundation (DEB-1518681; https://nsf.gov/). JRR was supported by the NSF (EF-1241889 and IOS-1754868; https://nsf.gov/), National Institutes of Health (R01GM109499 and R01TW010286-01; https://www.nih.gov/), and US Department of Agriculture (2009-35102-0543; https://www.usda.gov/wps/portal/usda/usdahome).

